# Long-term activity drives dendritic branch elaboration of a *C. elegans* sensory neuron

**DOI:** 10.1101/685339

**Authors:** Jesse A Cohn, Elizabeth R Cebul, Giulio Valperga, Mario de Bono, Maxwell G Heiman, Jonathan T Pierce

## Abstract

Neuronal activity often leads to alterations in gene expression and cellular architecture. The nematode *Caenorhabditis elegans*, owing to its compact translucent nervous system, is a powerful system in which to study conserved aspects of the development and plasticity of neuronal morphology. Here we focus on one sensory neuron in the worm, termed URX, which senses oxygen and signals tonically proportional to environmental oxygen. Previous studies have reported that URX has variable branched endings at its dendritic sensory tip. By controlling oxygen levels and analyzing mutants, we found that these branched endings grow over time as a consequence of neuronal activity. Furthermore, we observed that the branches contain microtubules, but do not appear to harbor the guanylyl cyclase GCY-35, a central component of the oxygen sensory transduction pathway. Interestingly, we found that although URX dendritic tips grow branches in response to long-term activity, the degree of branch elaboration does not correlate with oxygen sensitivity at the cellular or the behavioral level. Given the strengths of *C. elegans* as a model organism, URX may serve as a potent system for uncovering genes and mechanisms involved in activity-dependent morphological changes in neurons.

## INTRODUCTION

The nervous system often displays morphological plasticity in response to prolonged input or activity. These activity-dependent changes in neuron shape allow animals to interact more adeptly with their environment. For instance, the growth and pruning of specific synapses as well as axon and dendritic branches allow neural circuits to alter synaptic weighting during forms of learning and homeostatic plasticity [1, 2]. Interneurons also adjust the number and shape of their minute dendritic spines to filter input differently in neuronal networks [3–5]. In the sensory system, photoreceptor outer segment length has been shown to change in response to different light levels [6]. Thus, although the gross structure of the adult nervous system often remains static, many neurons change shape at subtle spatial and temporal scales.

The transparency, genetic tractability, and compact nervous system of the nematode *C. elegans* make the worm an excellent system to study genes that underlie how neurons achieve and adjust their shape. Many aspects of neuronal morphology have been examined in *C. elegans*, such as axonal and dendritic establishment [7, 8], dendritic tiling [9], synapse specification [10], and sensory cilia morphogenesis and maintenance [11, 12]. The worm has also been used to study how neurons alter their shape in response to changes in environment, such as the reshaping of the ciliated chemosensory neuron AWB by sensory activity [13], and the restructuring of sensory neuronal endings in an alternative developmental larval stage termed dauer, which is induced by certain environmental conditions [14–16]. Furthermore, many of the genes required for the development and maintenance of sensory cilia in *C. elegans* have conserved roles across species [17, 18].

Most sensory neurons in the head of *C. elegans* are bilaterally symmetric and have a cell body that projects a single dendrite to the tip of the nose, where the sensory transduction machinery is often localized [19]. Here we focus on one of these sensory neurons, the oxygen sensing neuron pair URX. In its natural environment where it burrows through rotting vegetation, *C. elegans* experiences a wide range of oxygen levels from nearly anaerobic patches (1% oxygen) to surface level oxygen (21%) [20]. When assayed in an oxygen gradient in the lab, worms exhibit a preference for 7-10% oxygen environments, which is thought to reflect the natural environment most suitable to their survival [21, 22]. This migratory behavior, termed aerotaxis, is primarily driven by the URX neurons. Mutant worms that lack components of the oxygen sensory transduction pathway in URX or worms that have URX ablated are deficient in aerotaxis [23, 24]. Calcium imaging experiments have revealed that, unlike many other sensory neurons in worm that respond phasically to changes in stimuli, URX neurons remain tonically active at surface levels of oxygen (21%) [25].

We report here that continuous exposure to surface level oxygen causes the URX neuron to steadily grow elaborate branches at its dendritic sensory ending over the course of adulthood. Branch elaboration depends on oxygen levels because cultivating worms in low oxygen (1% O_2_) prevented growth of these complex-shaped dendritic tips. We also find that the oxygen sensory pathway is necessary for this growth, suggesting that branch elaboration is due to neuronal activity. The components of the oxygen sensing pathway normally localize to a position at the end of the dendrite just beneath the surface of the nose of the worm, where they are thought to assemble into a signaling microdomain [26, 27]. Using a fluorescent tag, we examined the localization of one of the main components of the oxygen sensation pathway and found that the activity-dependent dendritic branches do not appear to be extensions of the oxygen signaling compartment. Consistent with this finding, we found that more elaborated dendritic branches do not increase sensitivity to oxygen at either the cellular or the behavioral level.

Sensory endings in *C. elegans* have been compared to several different structures in higher animals, such as dendritic spines, sensory cilia, and primary cilia [13, 28]. In addition, many of the components of the URX sensory cascade have homologous counterparts in the nervous systems of higher animals. URX may therefore serve as a powerful system for identifying important conserved genes and mechanisms involved in activity-driven morphological changes in other species.

## RESULTS

### Oxygen sensation drives dendritic branch elaboration in the oxygen-sensing neuron pair URX

*C. elegans* uses the bilaterally symmetric neuron pair URX to sense environmental oxygen levels [21, 22, 25]. The URX cell body is located in the head of the worm near the posterior pharyngeal bulb, and each URX neuron extends an axon into the nerve ring and a dendritic process to the nose of the worm (Figure 1A) [19, 29, 30]. This dendritic process ends just beneath the skin, where environmental oxygen may diffuse a short distance to bind the molecular receptor for oxygen, a class of membrane-anchored <u>g</u>uanylyl-<u>cy</u>clases (*gcy*) [26]. Previous studies of the *C. elegans* nervous system describe the URX dendritic tip as having branched endings that are variable in size and morphology [19, 29, 30]. While investigating gene expression in URX using fluorescence microscopy, we noticed that worms grown in low oxygen environments (1% O_2_) fail to sprout branches at the ends of the URX dendrites (Figure 1B left), in contrast to the branches seen in worms grown in high oxygen (21% O_2_) (Figure 1B right).

**Figure 1.**
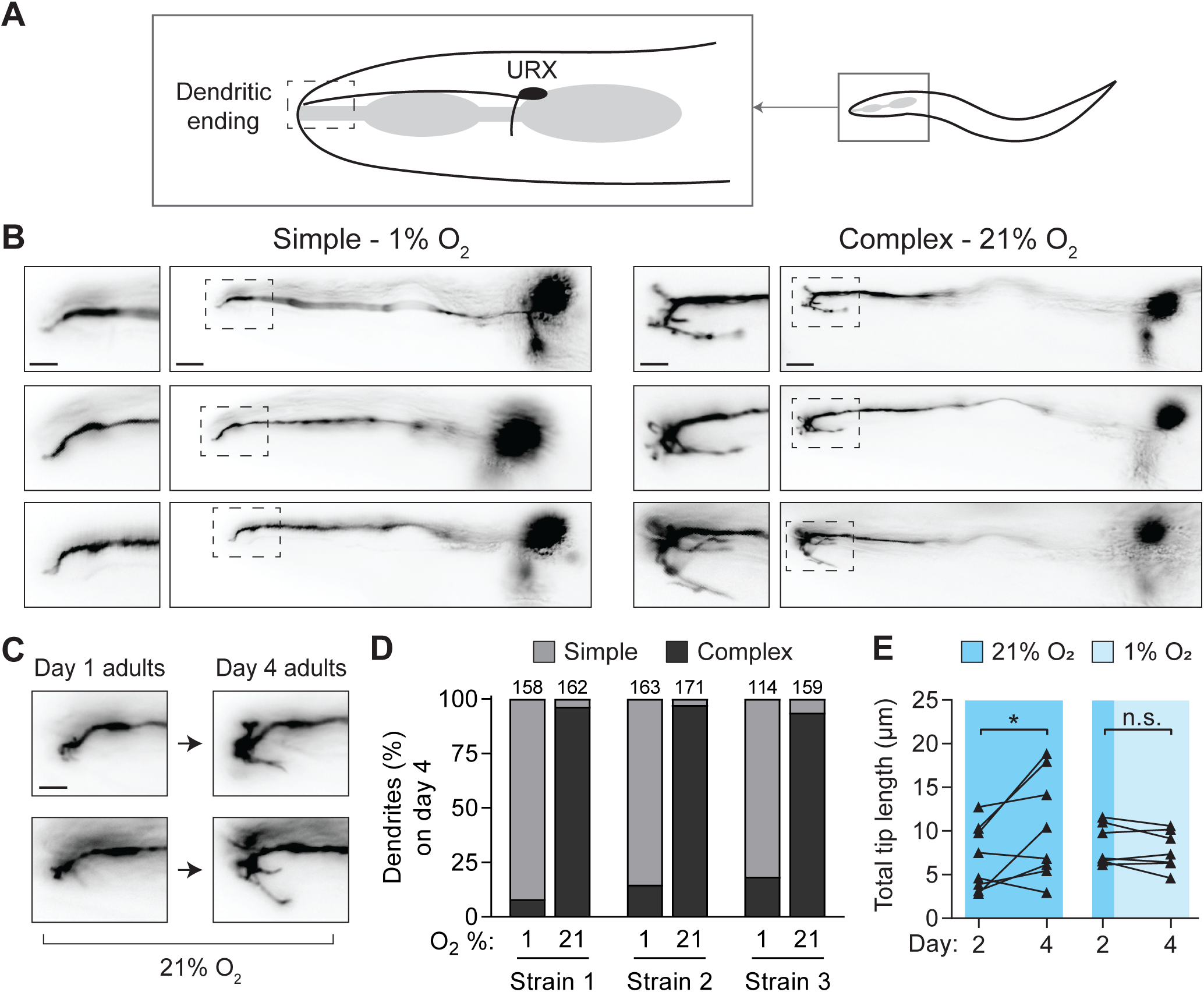
Environmental oxygen drives branching of the dendritic endings of the URX neuron pair. A.) Cartoon representation of the locations of the URX cell body and dendritic ending in the head of the worm. In all images, anterior is to the left, dorsal is upwards. B.) Examples of simple and complex dendritic ending morphologies in URX in wild-type worms. URX was visualized by expressing a *Pgcy-32::GFP* transgene, and worms were grown in either high (21%) or low (1%) oxygen until day four of adulthood. Note variable branched morphology of complex dendritic endings. Scale bars in images showing full neuron and inset are 10 µm and 5 µm, respectively. C.) The same individuals were imaged on day one and day four of adulthood to show the growth of branches over time in 21% oxygen. Scale bar is 5 µm. D.) Scoring of dendritic ending morphology in wild-type worms grown and maintained at either high or low oxygen until day four of adulthood. Three independently-derived strains were scored. In this and all similar figures, the number of dendrites scored per condition is shown above each bar. E.) Wild-type worms were reared in high oxygen until day two of adulthood, and then maintained in either high or low oxygen until day four of adulthood, with images taken at both time points. Branched endings in worms maintained at high oxygen showed significant growth over these two days (t = 2.5, p<0.05), while those in low oxygen neither grew nor shrank appreciably (t = 1.2, p>0.05). n = 8 for high oxygen conditions, 7 for low oxygen condition. Statistical significance determined by paired t-test.

To study the effect of oxygen level on branch elaboration, we visualized URX neurons using cytoplasmic GFP driven by the *gcy-32* promoter. This *gcy-32* reporter is robustly expressed in URX, AQR, and PQR neurons [31], but because neither AQR nor PQR send processes to the nose, we could clearly visualize the dendritic endings of URX in these strains. We characterized the dendritic morphology of URX in three independently-derived transgenic strains to control for artifacts caused by variation in GFP expression [32].

We imaged individual worms repeatedly across each day of adulthood and observed that URX dendritic branches in worms maintained at high oxygen levels continued to grow in length and complexity as the animal aged (Figure 1C). By day four of adulthood, the difference between the dendritic branches of worms grown in 21% and those grown in 1% was pronounced, so we chose this particular age to quantify differences between conditions. The morphological variability of the dendritic tips was difficult to describe; however, we found that we could unambiguously classify dendritic tips with elaborate branches as “complex”, and those without as “simple”. Specifically, if the dendritic tip had at least one secondary branch longer than 5 µm, we classified it as complex; otherwise the dendritic tip was classified as simple. In worms grown in 21% oxygen, the vast majority of dendritic tips were complex in each of the three transgenic reporter lines (complex = 96.3%, 97.1%, and 93.7%), while worms grown in 1% oxygen had mostly simple dendritic tips (complex = 8.2%, 14.7%, 18.4%) (Figure 1D). These results show that oxygen drives growth of elaborate branches at the end of the URX sensory dendrite, and that growth continues as long as the worm remains exposed to oxygen.

### Dendritic branches in URX are not actively broken down in low oxygen

The sensory transduction proteins in URX are localized to the dendritic ending [26], so we hypothesized that the growth of the complex branching might represent a form of homeostatic plasticity where the tips would expand in high oxygen and be broken down in low oxygen in order to regulate the amount of sensory receptors in the dendritic ending. To test this idea, we raised worms at 21% oxygen until day two of adulthood, at which point we quantified the total length of the branches on a single dendritic tip per worm. The imaged worms were then individually recovered and maintained for the next two days at either high or low oxygen, at which point we again quantified the total length of the branches (Fig 1E). We found that while the dendritic branches in worms kept in high oxygen (21%) continued to grow over the two days, the branches in worms moved to low oxygen (1%) neither grew nor reduced, but rather stayed the same length. Thus, dendritic branches in URX are not broken down in a low oxygen environment once established, and a visible “imprint” of high oxygen exposure remains encoded in the morphology of the URX ending.

### The oxygen sensing pathway is necessary for dendritic branch growth in URX

URX is a sensory neuron for oxygen, which suggests the possibility that branch growth at the dendritic tip is caused by prolonged sensory activity. In URX, molecular oxygen is coordinated at the dendritic tip by a heterodimer of the membrane-tethered guanylyl cyclases GCY-35 and GCY-36 [22, 26, 27]. Together these proteins produce intracellular cGMP when bound to oxygen, which in turn activates the cyclic nucleotide-gated cation channels TAX-4 and CNG-1 [22, 24, 33] (Figure 2A). We examined mutants lacking *gcy-35, cng-1,* or *tax-4*, and found that each mutant was profoundly defective in growing branches at the dendritic tip in URX when maintained at 21% oxygen until day four of adulthood (complex = 8.3%, 4.4%, 6.5%, respectively) (Figure 2B). This strongly suggests that oxygen sensation drives branch elaboration at the URX dendritic tip.

**Figure 2.**
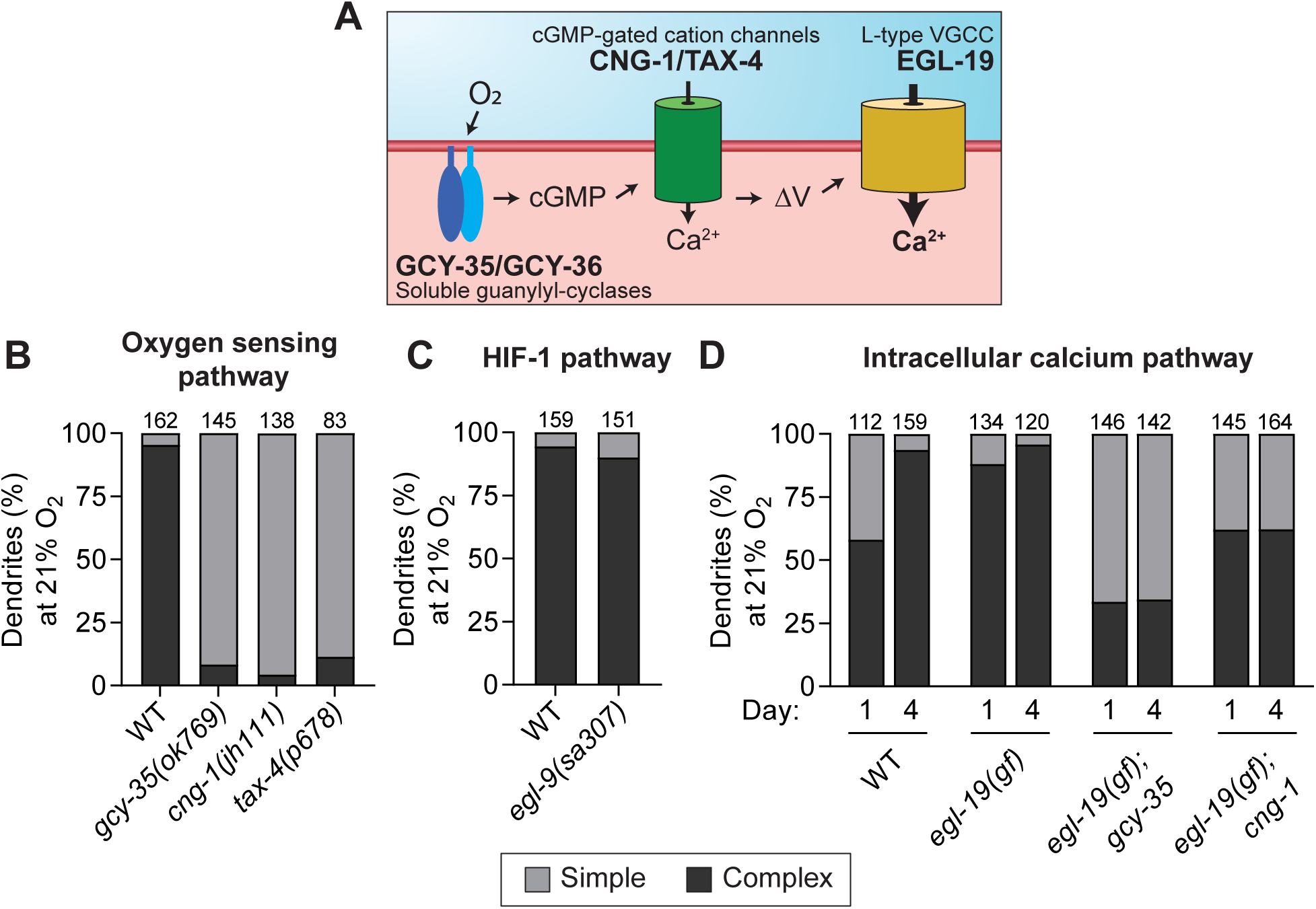
Sensory activity is necessary for elaboration of branched endings in URX. A.) Schematic showing the oxygen-sensing pathway in URX. A dendritically localized heterodimer of GCY-35/GCY-36 produces cGMP after binding oxygen. cGMP then activates downstream CNG cation channels CNG-1 and TAX-4. Membrane depolarization subsequently gates the L-type voltage-gated calcium channel EGL-19. B-C.) URX dendritic ending morphology was scored in worms of the given genotype on day four of adulthood after growth in high oxygen conditions. Components of the oxygen-sensing pathway were necessary for dendritic branch elaboration, while stabilization of HIF-1 in the *egl-9(sa307)* mutant had no effect on branch growth. Each bar is representative of three independently-derived transgenic strains expressing *Pgcy-32::GFP* to visualize URX. D.) URX dendritic ending morphology was scored in worms of the given genotype and age. Gain-of-function allele *egl-19(gf)* refers to the *egl-19(n2368)* allele.

We also considered the alternative hypothesis that branch growth is repressed by the hypoxia pathway in low oxygen (1%) and thus is revealed at high oxygen (21%). To test this hypothesis, we examined mutants lacking the prolyl hydroxylase EGL-9, which mediates degradation of the hypoxia pathway transcription factor HIF-1 under high oxygen conditions [34]. In *egl-9* mutants, the hypoxia pathway is constitutively active [35]. We found that the *egl-9* mutant had normal URX branching at 21% oxygen (complex = 90.1%, Figure 2C), indicating that HIF-1-dependent gene expression is likely not involved in repressing dendritic branch growth in low oxygen conditions.

Taken together, these results support the idea that the GCY-35/GCY-36-CNG-1-TAX-4 oxygen sensing pathway drives branch growth under high oxygen conditions, and that the lack of branch growth in low oxygen conditions is due to decreased activity of URX, not repression by the hypoxia pathway.

### Intracellular calcium influx is sufficient for dendritic branch growth

Both cGMP and calcium levels increase in URX as a result of oxygen sensation. To determine if calcium alone is sufficient for dendritic branch growth, we took advantage of a gain-of-function allele of *egl-19*, the sole L-type voltage-gated calcium channel in *C. elegans* [36]. The EGL-19 calcium channel is gated after depolarization of the URX neuron by CNG-1 and TAX-4 channels (Figure 2A). The *egl-19(n2368)* allele encodes an EGL-19 channel with an increased open probability compared to the wild-type protein, thus making it less dependent on CNG-1 and TAX-4 channel activity. This gain-of-function (*gf*) mutant background allowed us to test whether increased intracellular calcium influx in URX could induce branch growth in wild-type and the oxygen sensory transduction mutant backgrounds *gcy-35* and *cng-1* (Figure 2D).

First, we found that the *egl-19(gf)* allele in an otherwise wild-type background caused precocious branch elaboration in URX, with 88% of dendritic endings in day one adult worms showing complex branches, compared to only 58% in wild-type worms. This result suggests that extra calcium via EGL-19 may be sufficient to promote URX branch elaboration.

Second, we found that crossing the *egl-19(gf)* mutation to the *gcy-35* or *cng-1* oxygen sensory transduction mutants dramatically increased their portion of complex URX branches. We found that the URX dendritic ending in the *egl-19(gf);gcy-35* double mutant showed 35% complex branches, compared to 8.3% in the *gcy-35* single mutant. In the *egl-19(gf);cng-1* double mutant, 65% of URX dendritic endings had complex branches, compared to 4.4% in the *cng-1* single mutant. Intriguingly, we found that the URX dendritic ending in the *egl-19(gf);cng-1* double mutant often took on a blobby appearance when compared with other backgrounds, examples of which are shown in Supplemental Figure 1. The above results suggest that increased intracellular calcium influx via the EGL-19 L-type calcium channel is sufficient to stimulate elaboration of URX dendritic branches even in the absence of components of the oxygen sensation cascade.

### Wild-type URX dendrites have exuberant branching and an intricate internal structure

We next used super-resolution microscopy to gain a more detailed look at the dendritic branches or lack thereof in wild-type and *gcy-35(ok769)* day-four adult worms (Figure 3). These higher resolution pictures of URX confirm the complex morphology of dendritic endings in wild-type worms and the lack of branched dendritic endings in the *gcy-35* mutant. In addition, we noticed several interesting features at this resolution that we were unable to discern by conventional microscopy. For one, nearly all *gcy-35* mutant individuals had a small membranous fan structure attached to the end of the dendritic stalk. This is reminiscent of the membranous fan-like structures that were observed in the dendritic tips of the AWB chemosensory neurons in similar sensory transduction mutants [13, 29, 30]. We also noticed several areas in the dendritic stalk and in the outgrown branches that excluded GFP, leading to an intricate and complicated internal structure. These exclusions are likely secretory vesicles, and may reflect dense secretory traffic in the wild-type dendrite to facilitate branching at the dendritic ending..

**Figure 3.**
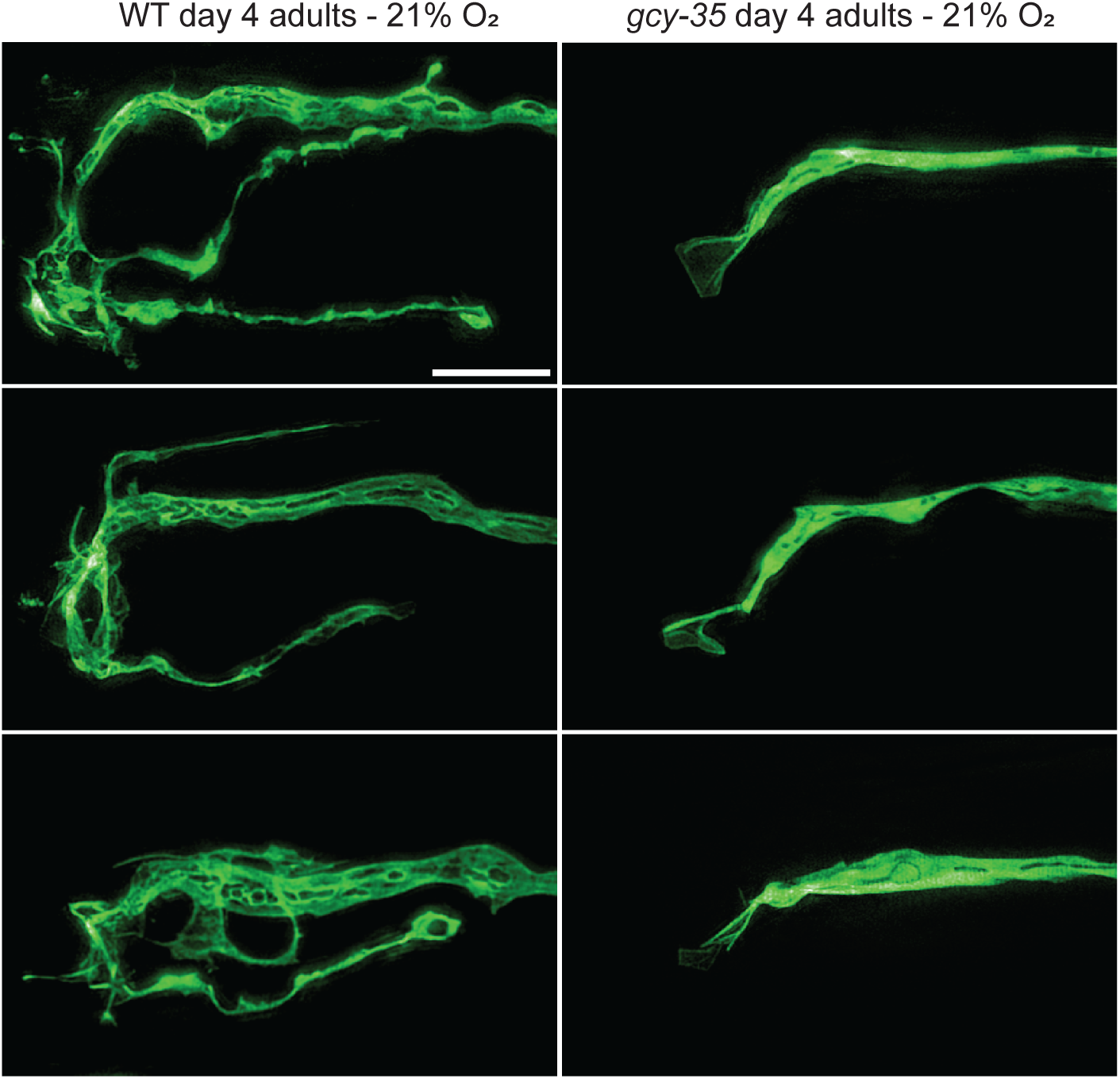
High-resolution images of branched URX dendritic endings. Representative high-resolution images of the dendritic endings of URX in either wild-type or *gcy-35(ok769)* mutant worms. All animals were grown in high oxygen until day four of adulthood. Scale bar is 5 µm.

### Branched URX dendritic endings contain microtubules and do not appear to expand an oxygen sensory compartment

To learn more about the nature of the branched dendritic endings in URX, we utilized a strain in which the microtubule binding protein EBP-2 is tagged with GFP in order to determine whether the cellular cytoskeleton extends into the outgrown dendritic branches [37]. In day four adult worms, we found that EBP-2::GFP was present throughout URX, including the dendritic branches, in all individuals examined (n = 15/15). This suggests that microtubules may play a role in cytoskeletal support and/or cellular transport in these branched dendritic endings.

Components of the oxygen-sensing machinery, including the guanylyl cyclases GCY-35 and GCY-36, localize to the ending of the dendritic stalk in URX, where they are thought to associate with one another to form a signaling microdomain [26, 27]. We used a strain in which GCY-35 is tagged with GFP to investigate whether the dendritic branches in day four adults contain components of the oxygen sensing pathway and may therefore be an expansion of the URX oxygen sensory compartment (Figure 4B). We found that in all worms examined, GCY-35::GFP localized to the end of the dendritic stalk in the nose, and in 75% of worms (n = 63/84), GCY-35::GFP was not visible in the outgrown dendritic branches. In the remaining 25% of worms (n = 21/84), we noticed GFP signal in the outgrown branches, though usually at a much lower level than is seen at the end of the dendritic stalk. Although it is possible that GCY-35::GFP is present at levels below our limit of detection, or untagged GCY-35 is present in the branches, we interpret these data to mean that the outgrown branches at the dendritic ending of URX are likely not an expansion of the sensory compartment.

**Figure 4.**
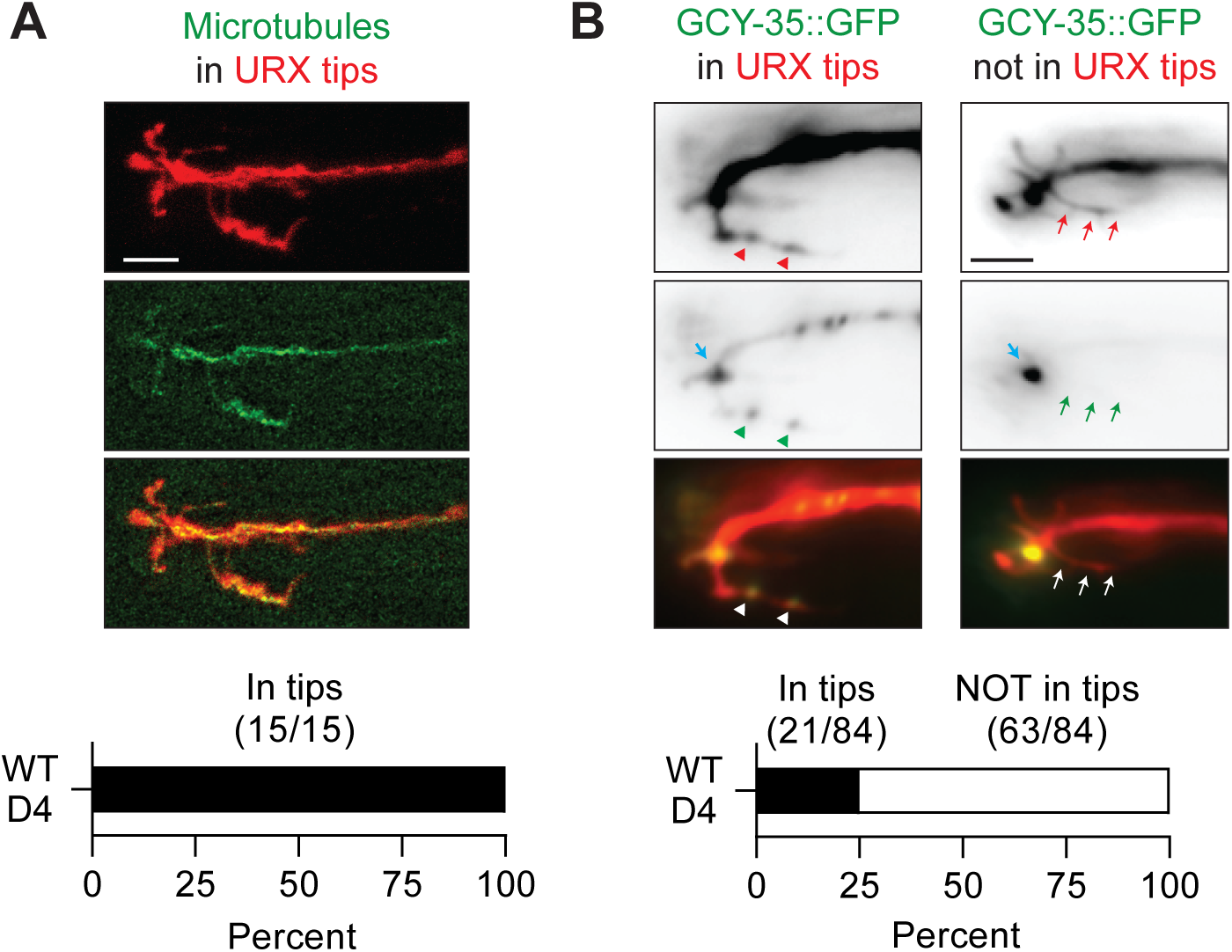
Branched URX dendritic endings contain microtubules and are likely not an expansion of an oxygen sensory compartment. A.) An example day four wild-type adult grown in high oxygen and expressing soluble mCherry cell-specifically to label URX, and EBP-2::GFP in URX to label microtubules. Scale bar is 5 µm. B.) Localization of GCY-35::GFP in URX in day four wild-type adults grown in high oxygen. Blue arrows label a likely sensory compartment in URX where GCY-35::GFP is highly concentrated in all animals. 25% of animals examined had visible GFP signal in the outgrown dendritic branches (left), while 75% did not (right). Scale bar is 5 µm.

### Dendritic branches do not overtly contribute to cellular oxygen sensation or an oxygen-related behavior

URX responds to oxygen tonically. Ambient environmental oxygen causes a proportional steady intracellular calcium influx into URX [25]. This in turn causes URX to signal downstream interneurons and thereby drive oxygen-related behaviors. Because some sensory neurons have distinct spatial compartments that contribute to different aspects of their function [38], we hypothesized that the complex branches at the URX dendritic tip might enhance oxygen sensing. If so, enhanced oxygen sensation may be reflected by alterations in cellular calcium responses to oxygen or in oxygen-related behaviors. To test this hypothesis, we chose to compare the cellular and behavioral responses to oxygen in day one adults and day four adults grown in 21% oxygen, because day four adults have more elaborated and developed branches at the URX dendritic ending than day one adults. We chose not to study the cellular and behavioral responses to oxygen for the sensory mutants described above because although they fail to grow complex branches, they are impaired in oxygen sensation. Likewise, we did not study wild-type worms maintained in 1% oxygen because although this prevents complex branching, it may also cause the neuron to adapt in a way that would confound our results [26].

To compare the cellular response to oxygen, we used the ratiometric calcium indicator yellow cameleon 2.60 (YC2.60) to measure calcium levels in the cell body of URX in response to step changes in environmental oxygen from 7% O_2_ to 21% O_2_ and back to 7% O_2_. We found that day one adults and day four adults displayed indistinguishable calcium responses in this paradigm. In both cases, calcium levels appeared to increase and stay sustained to a similar level and return to a similar baseline with the same time course (Figure 5A). These results suggest that differences in dendritic branch complexity and length do not correlate with differences in the amplitude of calcium influx in the URX cell body in response to oxygen, fitting with our finding above that the branches likely do not represent expansions of the URX oxygen sensory compartment.

**Figure 5.**
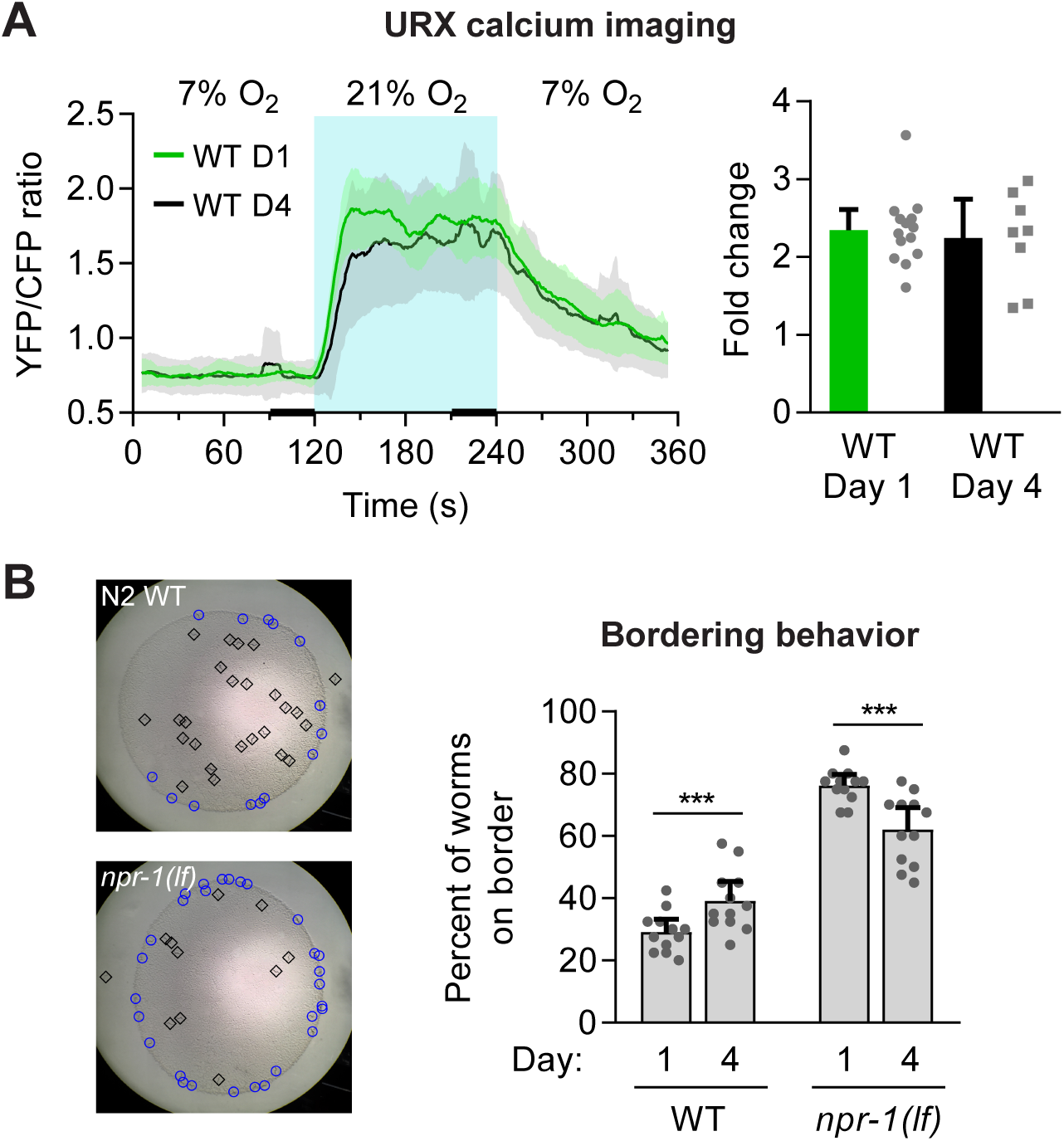
URX dendritic ending branch length/complexity does not correlate with increased oxygen sensitivity in URX. A.) Wild-type URX calcium responses to 7% −21% −7% oxygen steps. No significant difference in baseline calcium levels or response size was seen across days of adulthood. Shaded areas are 95% confidence intervals. Thick black lines near x-axis show the time intervals used to calculate fold change. N=14 for day one adults, 8 for day four adults. Error bars on fold change are 95% C.I. B.) Left images show examples of the URX-dependent bordering behavior in wild-type and *npr-1(ky13)* day one adults. To the right, bordering behavior in wild-type or *npr-1(ky13)* worms was quantified on day one or day four of adulthood. Opposing trends suggests that URX dendritic branch complexity does not contribute to this behavior. Each point is an assay of 40 worms. Error bars are 95% C.I., statistical significance determined by Student’s t-test, *** indicates p < 0.01.

We also examined a URX-dependent behavior called bordering. Wild isolates of *C. elegans* prefer conditions of 5-12% environmental O_2_ and avoid higher levels of oxygen [22]. Consequently, wild isolate worms in the lab will accumulate at the border of a bacterial lawn where the bacteria grows thickest and local oxygen levels are closest to their preferred level. Worms lacking URX or components of the oxygen sensory pathway no longer display this bordering behavior [21, 22]. A gain-of-function mutation in the gene *npr-1* in the standard lab strain N2 causes them to have a blunted oxygen aversion compared to wild isolates, and therefore N2 worms also have a less pronounced bordering behavior [39]. N2 worms with a loss-of-function in *npr-1* border similarly to true wild isolates, and also grow branches at the URX dendritic ending like the standard N2 strain (Supplemental Figure 2).

We hypothesized that if the complex branches of URX dendritic tips enhanced oxygen sensation in a manner that was missed by our cellular imaging experiment above, perhaps the oxygen-dependent bordering behavior would be increased in older adults compared to young adults. We compared day one and day four worms of either N2 or *npr-1* loss-of-function backgrounds. Oddly, we found opposing results in these two backgrounds – that is, day four N2 worms bordered slightly more strongly than day one N2 worms, while day four *npr-1(lf)* worms bordered less strongly than day one *npr-1(lf)* worms (Figure 5B). The effect sizes of these differences were small but statistically significant. We interpret these unexpected opposing results to suggest that dendritic branch length/complexity does not correlate with a difference in bordering behavior at these two ages. This is in agreement with our above finding that the URX cellular response to oxygen does not correlate with dendritic branch complexity. Taken together, these sets of experiments indicate that the growth of branched dendritic endings correlates with but does not contribute to oxygen sensation in URX.

## DISCUSSION

### Morphological changes in *C. elegans* sensory neurons

*C. elegans* is a powerful system for studying changes in neuron morphology, as the transparency of the animal allows for convenient imaging of fluorescently labeled, identified cells in live animals across different ages and conditions. The *C. elegans* literature has several examples of developmental changes in neuronal shape, such as the pruning of excessive neurites [40], and the restructuring of certain sensory neurons during the dauer alternate larval stage [14–16]. There are also a few examples in adulthood, such as the age-dependent increase in the branching of the PVD mechanosensory neurons [41], and the abnormal morphology of some sensory neuron endings in signal transduction mutant backgrounds [42, 43]. However, to our knowledge, there is only one other clear example of environmental input directly influencing sensory ending morphology in *C. elegans*, which is that of the AWB chemosensory neurons [13]. The AWB sensory dendritic ending has been shown to remodel from two finger-like ciliated branches to two fan-like structures in the absence of olfactory sensory input. This change was dependent on sensory transduction genes and the kinesin-II motor protein, among factors. Interestingly, several other neurons in *C. elegans* with characteristic morphologies, such as AFD, do not show obvious remodeling of their sensory ending depending on neuronal activity [42]. Why some sensory neurons in *C. elegans* undergo shape changes in response to activity while others do not is an intriguing question for further analysis.

Recently, McLachlan et al. described a role for the *C. elegans* MAPK15/ERK8 homologue in controlling dendritic length in URX [44]. In the *mapk-15* mutant, URX grows extremely long dendritic endings. Despite these long dendritic overgrowths, the *mapk-15* mutant displayed a roughly normal response to oxygen at both cellular and behavioral levels. This mirrors our finding that the presence or absence of branched endings had no discernible effect on cellular physiology or behavior. One interesting difference between the dendritic overgrowths in the *mapk-15* mutant and the shorter branches we see in wild-type URX neurons is the localization of the GCY-35 guanylyl-cyclase. In the *mapk-15* mutant, GFP-tagged GCY-35 was observed throughout the dendritic overgrowths, whereas we did not see GCY-35::GFP in the complex dendritic branches of most wild-type worms. One explanation for this discrepancy might be that MAPK-15 defines and constrains the size of an oxygen-sensing compartment at the end of the URX sensory dendrite. In the *mapk-15* mutant then, this sensory compartment is expanded, whereas in wild type, the complex dendritic outgrowths we see represent a distinct subcellular compartment. Though this idea is speculative, sensory ending compartmentalization has been reported in other *C. elegans* neurons [38].

### URX dendritic branch function

In our assays, longer and more complex branches at the URX dendritic tip did not correlate with detectable differences in oxygen sensation as measured at both the cellular and the behavioral level. It remains to be seen then what, if any, consequences branching of the URX dendritic ending has on physiology or behavior in *C. elegans*. We offer a few hypotheses, though further experimentation is necessary to test these ideas.

For one, because dendritic outgrowths would increase the volume of the dendritic ending, this may dilute local secondary messengers including cGMP. At 21% oxygen cGMP is actively and continuously produced in URX [25, 33]. If degradation of cGMP by cellular phosphodiesterases does not completely balance production of cGMP by guanylyl-cyclases, this dilution may represent a homeostatic mechanism by which the cell maintains steady calcium influx levels. Similarly, the increase in plasma membrane could lead to an increased capacitance at the sensory ending, balancing prolonged cation influx to normalize downstream voltage-gated signaling. In these scenarios, we might not expect to see differences in calcium influx at the cell body and/or in behavior between day one and day four adults, because each of these mechanisms serves to normalize the signal that reaches the cell body.

Alternatively, the complex dendritic outgrowths may contain sensory transduction molecules other than those already known for oxygen sensation. In this study, we only examined the localization of the oxygen sensor guanylyl-cyclase GCY-35 and found that in most animals it localized only in its expected position at the anterior tip of the dendritic stalk, rather than in the outgrown branches. Because branch length and complexity did not correlate with oxygen responses, this raises the intriguing possibility that URX may also sense another stimulus, perhaps one that correlates with surface level oxygen in the natural environment of *C. elegans*. Although to this point URX is only known to sense oxygen, several sensory neurons in *C. elegans* are polymodal [45–49]. In regard to this idea, it is notable that URX uniquely expresses several guanylyl-cyclases with currently unknown functions including GCY-25 and GCY-37 [31]. Although these ideas are only conjecture, it will be interesting to try to deduce a physiological role for branched sensory endings in URX in future studies.

In summary, we have shown that the URX sensory neurons in *C. elegans* grow an elaborate branched structure at their dendritic endings in an activity-dependent manner. In all animals, the ability for neurons to alter aspects of their shape is central to their proper functioning. *C. elegans* has contributed much to our understanding of the genes and rules involved in the organization and morphogenesis of the nervous system in mammals; thus URX may serve as a powerful system for identifying genes important in activity-dependent shape changes in neurons in higher animals.

## ACKNOWLEDGMENTS

We would like to thank Martin Harterink and Casper Hoogenraad for sharing strains; Edward Mills for the oxygen chamber; Anna Kazatskaya and Piali Sengupta for sharing unpublished results; Talley Lambert and Jennifer Waters (Harvard Medical School Cell Biology Microscopy Facility) for their assistance with SIM acquisition; and Lin Shao (Yale University) for CUDA-accelerated 3D-SIM reconstruction code. Some strains were provided by the CGC, which is funded by NIH Office of Research Infrastructure Programs (P40 OD010440). Additional funds were provided by NIH F31NS103371 and NIH-NIA grants RF1AG057355 and R01AG041135.

## MATERIALS AND METHODS

### Strains

The following strains were used:

N2 Bristol; JPS879 *vxEx879[Pgcy-32::GFP Punc-122::GFP]*; JPS880 *vxEx880[Pgcy-32::GFP Punc-122::GFP]*; JPS881 *vxEx881[Pgcy-32::GFP Punc-122::GFP]*; JPS882 *gcy-35(ok769) I; vxEx882[Pgcy-32::GFP Punc-122::GFP]*; JPS883 *gcy-35(ok769) I; vxEx883[Pgcy-32::GFP Punc-122::GFP]*; JPS884 *gcy-35(ok769) I; vxEx884[Pgcy-32::GFP Punc-122::GFP]*; JPS941 *cng-1(jh111) V; vxEx941[Pgcy-32::GFP Punc-122::GFP]*; JPS942 *cng-1(jh111) V; vxEx942[Pgcy-32::GFP Punc-122::GFP]*; JPS943 *cng-1(jh111) V; vxEx943[Pgcy-32::GFP Punc-122::GFP]*; JPS921 *tax-4(p678) III; vxEx921[Pgcy-32::GFP Punc-122::GFP]*; JPS922 *tax-4(p678) III; vxEx922[Pgcy-32::GFP Punc-122::GFP]*; JPS923 *tax-4(p678) III; vxEx923[Pgcy-32::GFP Punc-122::GFP]*; JPS1103 *egl-9(sa307) V; vxEx1103[Pgcy-32::GFP Punc-122::GFP]*; JPS1104 *egl-9(sa307) V; vxEx1104[Pgcy-32::GFP Punc-122::GFP]*; JPS1105 *egl-9(sa307) V; vxEx1105[Pgcy-32::GFP Punc-122::GFP]*; JPS1079 *egl-19(n2368) IV; vxEx1079[Pgcy-32::GFP Punc-122::GFP]*; JPS1159 *gcy-35(ok769) I; egl-19(n2368) IV; vxEx1079[Pgcy-32::GFP Punc-122::GFP]*; JPS1160 *egl-19(n2368) IV; cng-1(jh111) V; vxEx1079[Pgcy-32::GFP Punc-122::GFP]*; JPS1161 *hrtSi4[Pgcy-36::EBP-2::GFP]; vxEx1161[Pgcy-32::mCherry +rol-6(su1006)]*; JPS1123 *dbEx[Pgcy-37::GCY-35::HA::GFP::SL2::mCherry]*; AX7313 *vxIs601[Pegl-1::mCherry Punc-122::GFP]; dbEx613[Pgcy-37::YC2.60 Punc-122::RFP]*; CX4148 *npr-1(ky13) X*; JPS1173 *npr-1(ky13) X; vxEx1173[Pgcy-32::GFP Punc-122::GFP]*; JPS1174 *npr-1(ky13) X; vxEx1174[Pgcy-32::GFP Punc-122::GFP]*; JPS1175 *npr-1(ky13) X; vxEx1175[Pgcy-32::GFP Punc-122::GFP]*

### Microscopy and Dendrite Scoring

For the dendritic scoring assays, worms were synchronized by timed egg laying and then maintained either on the benchtop for 21% oxygen conditions, or in a Modular Incubator Chamber (Billups-Rothenberg) attached to an oxygen tank containing 1% O_2_ balanced with nitrogen (Airgas). On the day of the assay, either day one or day four adults were mounted on 2% agarose pads, anesthetized with 30-mM sodium azide diluted in NGM, then visualized on an Olympus IX51 inverted microscope equipped with an X-Cite FIRE LED Illuminator (Excelitas Technologies Corp.) and an Olympus UPlanFL N 40X/0.75 NA objective. Epifluorescence images were taken with a Retiga 2000R CCD camera (QImaging) and QCapture Pro 6.0 software.

Dendrites were scored as complex if they had at least one secondary branch that extended ≥5 µm from the primary dendritic stalk ending, and were scored as simple otherwise. We found that in some worms one of the URX neurons would be simple while the other would be complex, and a small number of worms expressed the *Pgcy-32::GFP* transgene in only one neuron of the URX pair. Therefore we scored each dendrite individually, giving an n of 2 or 1 per animal. Image analysis was performed using ImageJ (NIH). For Figure 1D, dendritic branch length was quantified using the segmented line tool in ImageJ. Pictures were analyzed in pairs to ensure consistent start and end points for branch measurements but blinded to age and condition.

Confocal images were captured with a Zeiss LSM 710 microscope equipped with a Plan-Apo 63X (oil); 1.4 NA objective lens and Zen Imaging software. Images were analyzed in ImageJ and are shown as maximum intensity Z-projections.

For SIM imaging, day four adults grown at 20°C were anesthetized in 110mM sodium azide and 20mM levamisole in M9, then mounted with No. 1.5 coverslips on 3% agarose pads with 110 mM sodium azide. Imaging was performed on an OMX V4 Blaze microscope (GE Healthcare) equipped with three watercooled PCO.edge sCMOS cameras, a 488 nm laser, and a 528/48 emission filter (Omega Optical). Images were acquired with a 60X/1.42 NA Plan Apochromat objective (Olympus) and a final pixel size of 80 nm. Z-stacks of ∼3-6 µm were acquired with a z-step of 125 nm and with 15 raw images per plane (three angles with five phases each). Spherical aberration was minimized using immersion oil matching [50]; generally, oil with a refractive index of 1.524 worked well. Superresolution images were computationally reconstructed from the raw data sets with a channel-specific, measured optical transfer function, and a Wiener filter constant of 0.001 using custom written 3D-SIM reconstruction code (T. Lambert, Harvard Medical School) based on Gustafsson et al. [51]. Images are displayed as maximum intensity Z-projections. Final image layouts were assembled in Adobe Illustrator.

### Molecular Biology and Transgenic Strain Construction

Constructs to label URX consisted of an 876-bp fragment of the *gcy-32* promoter amplified from genomic DNA, and either worm-optimized GFP amplified from plasmid pPD95.75, or worm-optimized mCherry amplified from plasmid pCFJ90. These fragments were combined by fusion PCR and then subcloned into a pCR-Blunt vector with the Zero Blunt PCR Cloning Kit from ThermoFisher Scientific. Each construct was injected at 20ng/µl along with either *Punc-122::GFP* or pRF4/*rol-6(su1006)* as a co-injection marker.

Mutations were followed in crosses by either PCR genotyping and additionally by phenotype when possible.

### Calcium Imaging

To image day one adults, we picked L4 animals expressing the YC2.60 Ca^2+^ sensor 24 h before imaging. To image day four adults, we picked L4s five days before the assay. On the day of the assay 5 – 10 worms were glued to agarose pads (2% in M9 buffer, 1 mM CaCl_2_), using Dermabond tissue adhesive, with their body immersed in OP50 washed off from a seeded plate using M9 buffer. The animals were quickly covered with a PDMS microfluidic chamber and 7% O_2_ pumped into the chamber for 2 min before we began imaging, to allow animals to adjust to the new conditions. Neural activity was recorded for 6 minutes with switches in O_2_ concentration every 2 minutes.

Imaging was on an AZ100 microscope (Nikon) bearing a TwinCam adaptor (Cairn Research, UK), two ORCA-Flash4.0 V2 digital cameras (Hamamatsu, Japan), and an AZ Plan Fluor 2× lens with 2× zoom. Recordings were at 2Hz. Excitation light from a Lambda LS xenon arc lamp (Sutter) was passed through a 438/24 nm filter and an FF458-DiO2 dichroic (Semrock). Emitted light was passed to a DC/T510LPXRXT-Uf2 dichroic filter in the TwinCam adaptor cube and then filtered using a 483/32 nm filter (CFP), or 542/27 nm filter (YFP) before collection on the cameras. Recordings were analysed using Neuron Analyser, a custom-written Matlab program available at https://github.com/neuronanalyser/neuronanalyser.

### Bordering Assays

Bordering assays were performed as previously described [26]. Worms were synchronized by selecting L4-stage hermaphrodites and letting them grow to either day one or day four adults for the assay. For each assay plate, 40 adult worms of the appropriate age were transferred to 6-cm diameter NGM plates that had been seeded two days prior with 200 µl of an overnight OP50 strain bacterial culture. One hour after transfer, the percentage of worms on the bacterial lawn border was calculated.

---

**Supplemental Figure 1. Example images of simple dendritic endings in *egl-19(gf)* double mutants.** Representative images of simple morphology in *egl-19(gf);gcy-35* day 4 adults and *egl-19(gf);cng-1* day four adults, all grown in high oxygen. The *egl-19(gf);gcy-35* double mutant had normal simple endings, while the *egl-19(gf);cng-1* double mutant had abnormal “blobby” endings. Scale bars in images showing full neuron and inset are 10 µm and 5 µm, respectively.

**Supplemental Figure 2.** T**h**e ***npr-1* loss-of-function mutant have normal branched dendritic URX endings.** Quantification of dendritic ending morphology in day four wild-type and *npr-1(ky13)* mutant worms grown in high oxygen.

